# *G*α*_olf_* Regulates Biochemical Signaling in Neurons Associated with Movement Control and Initiation

**DOI:** 10.1101/2024.04.03.587766

**Authors:** Michael Millett, Anika Heuberger, Elisabeth Martin Castosa, Allison Comite, Preston Wagner, Dominic Hall, Ignacio Gallardo, Nicole E Chambers, Lloyd Wagner, Mark S Moehle

## Abstract

The heterotrimeric G-protein α subunit, Gα_olf_, acts to transduce extracellular signals through G-protein coupled receptors (GPCRs) and stimulates adenylyl cyclase mediated production of the second messenger cyclic adenosine monophosphate. Numerous mutations in the *GNAL* gene, which encodes Gα_olf_, have been identified as causative for an adult-onset dystonia. These mutations disrupt GPCR signaling cascades in *in vitro* assays through several mechanisms, and this disrupted signaling is hypothesized to lead to dystonic motor symptoms in patients. However, the cells and circuits that mutations in *GNAL* corrupt are not well understood. Published patterns of Gα_olf_ expression outside the context of the striatum are sparse, conflicting, often lack cell type specificity, and may be confounded by expression of the close *GNAL* homolog of *GNAS*. Here, we use RNAScope in-situ hybridization to quantitatively characterize *Gnal* mRNA expression in brain tissue from wildtype C57BL/6J adult mice. We observed widespread expression of *Gnal* puncta throughout the brain, suggesting Gα_olf_ is expressed in more brain structures and neuron types than previously accounted for. We quantify transcripts at a single cell level, and use neuron type specific markers to further classify and understand patterns of *GNAL* expression. Our data suggests that brain regions classically associated with motor control, initiation, and regulation show the highest expression of *GNAL*, with Purkinje Cells of the cerebellum showing the highest expression of any neuron type examined. Subsequent conditional *Gnal* knockout in Purkinje cells led to markedly decreased intracellular cAMP levels and downstream cAMP-dependent enzyme activation. Our work provides a detailed characterization of *Gnal* expression throughout the brain and the biochemical consequences of loss of Gα_olf_ signaling *in vivo* in neurons that highly express *Gnal*.

## Introduction

The *GNAL* gene, located on chromosome 18p11, contains 12 exons that encode for heterotrimeric G-protein α subunit Gα_olf_ (Overhauser et al., 1993 Vuoristo et al., 2000). While originally characterized in olfactory epithelium, expression and activity of Gα_olf_has been noted in several brain structures (Jones & Reed, 1989). However, there has been a unique focus on the activity of Gα_olf_ in the striatum, where Gα_olf_ is classically thought to “replace” Gα_s_ as the primary source of stimulatory G protein signaling. In striatal spiny projection neurons (SPNs), Gα_olf_associates with Gβ_2_γ_7_ subunits to form a heterotrimer (Schwindinger et al., 2010a; Watson et al., 1994). Notably, the Gα_olf_β_2_γ_7_heterotrimer functions to transmit signal from D1 dopamine receptors (D_1_Rs) in the basal ganglia direct pathway and adenosine 2A receptors (A_2a_Rs) in the basal ganglia indirect pathway (Herve et al., 1993; Kull et al., 2000). Biochemically, Gα_olf_appears to act in a similar manner to its close homolog Gα_S_. Upon receptor activation, the Gα_olf_subunit undergoes conformational change, exchanging GDP for GTP, and dissociates from Gβ_2_γ_7_ dimer. After dissociation, active Gα_olf_ stimulates adenylyl cyclase isotype 5 (AC5) conversion of adenosine triphosphate (ATP) into cyclic adenosine monophosphate (cAMP)(Glatt & Snyder, 1993) Cooper, 2003). *GNAL* expression and Gα_olf_ activity outside the striatum remain largely undefined.

Multiple autosomal dominant *GNAL* mutations have now been identified as causative for primary dystonia (Fuchs et al., 2013; Kumar et al., 2014). These mutations lead to an adult onset, isolated dystonia that is largely indistinguishable from idiopathic dystonia. Utilizing *ex vivo* or *in vitro* expression systems, dystonia linked *GNAL* mutations have now been shown to cause abnormal Gα_olf_ signal transduction and subsequent downstream signaling through a number of mechanisms (Masuho et al., 2018). As Gα_olf_is the primary source of stimulatory G protein signaling in striatal SPNs, pathogenic Gα_olf_ signaling in the striatum is hypothesized to underlie dystonic motor symptoms in patients (Fuchs et al., 2013; Kumar et al., 2014; Masuho et al., 2018).

Rodent models are commonly employed by researchers to study the molecular and motor deficits of inherited dystonia (Imbriani et al., 2020). Expression of *Gnal* has been described in the cortex, striatum, hippocampus, thalamus, hypothalamus, medial habenula, amygdala, cerebellum, and midbrain via in-situ hybridization (ISH), northern blotting, sequencing, and immunohistochemical staining (Belluscio et al., 1998; Herve et al., 1993, 2001; Vemula et al., 2013; Vuoristo et al., 2000). Nevertheless, *GNAL*-linked dystonia research has classically focused on the striatum where pathogenic Gα_olf_ signaling has been observed in multiple systems. Despite recent evidence implicating cerebellar dysfunction resulting from *Gnal* loss in dystonia pathophysiology (Aïssa et al., 2022), the extent to which *Gnal* is expressed and the significance of Gα_olf_ signaling in most brain structures is unexplored.

Here, we use RNAScope, a sensitive in-situ hybridization technique, that allows for quantification of single mRNA transcripts, to characterize the expression of *Gnal* throughout the brain of C57BL/6 mice. We co-probe for markers of various neuron types to count the number of *Gnal* transcripts within and quantitatively compare expression between neurons of interest. Furthermore, we delineate relative expression levels of mRNA encoding for the predominant stimulatory G-protein α subunit expressed in the brain, Gα_s_ (Gilman, 1987). Finally, based on our quantification, we selectively remove *Gnal* in the cells that express the highest level of *Gnal*, Purkinje Cells of the cerebellum, to understand how loss of *Gnal* effects signaling in these cells. Our results help to identify neuron types and circuits to understand how pathogenic Gα_olf_ signaling leads to dystonia.

## Results

### *Gnal* is Widely Expressed in Murine Brain Tissue

To understand broadly where *Gnal* is expressed in the brain, we began by looking at *Gnal* mRNA transcripts using RNAscope mRNA ISH probes specific for *Gnal* (all reagents listed in Table 1). Expression of *Gnal* mRNA was observed throughout the majority of brain structures of adult C57Bl/6J mice, with apparent highest expression in cortex, striatum, hippocampus, substantia nigra, and cerebellum (Fig. 1). The translation of *Gnal* transcripts into protein was confirmed with Gα_olf_ antibody labeling (Sup. Fig. 1). RNAscope 3-plex positive and negative controls were used to optimize pretreatment conditions and determine proper acquisition settings, and show specificity of signal from ISH probes (Sup. Fig. 2).

**Figure 1:**
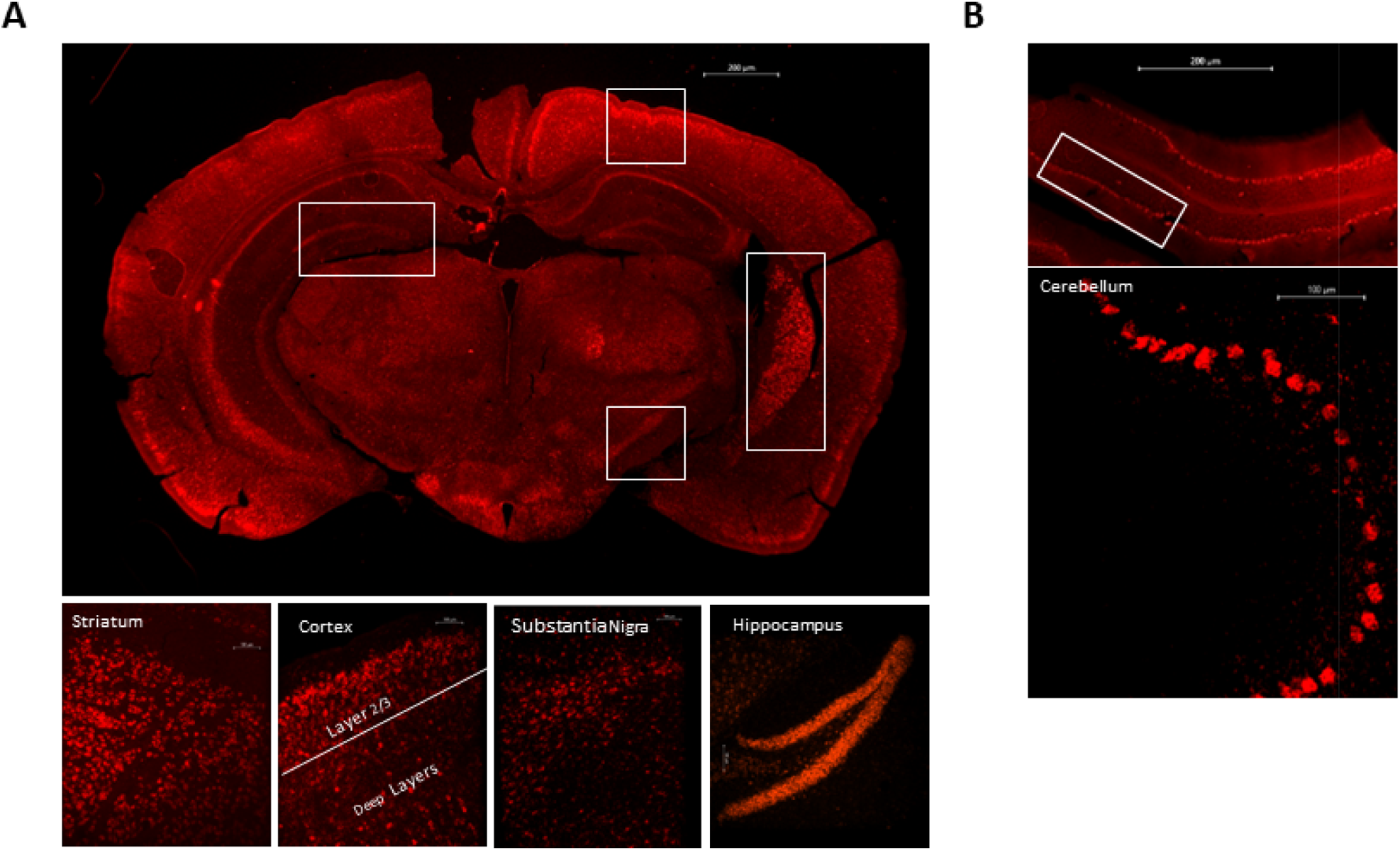
*Gnal* is Widely Expressed in Brain Tissue. A) Micrograph of *Gnal* (red) transcript expression in a representative coronal section of adult C57BL/6J brain tissue (top image). White boxes identify regions of concentrated labeling. Bottom row are enlarged representative pictures of each boxed region. B) Micrograph of *Gnal* transcript expression (red) patterns in an individual cerebellar branch of a hemisected sagittal section of C57BL/6J brain tissue (top image). White box highlights Purkinje cell layer. A representative pattern of *Gnal* expression is enlarged in bottom image. N= 3 mice.

**Table 1.**

| RNAScope Reagents: |
| --- |
| 1. RNAscope Multiplex Fluorescent Detection Reagents v2 (Cat. No. 323110) |
| 2. RNAscope™ Probe- Mm-Gnal (ACD Cat No. 411071) |
| 3. RNAscope™ Probe- Mm-Gnal-C2 ( ACD Cat No. 411071-C2) |
| 4. RNAscope™ Probe- Mm-Gnas-O1-C2 (ACD Cat No. 526891-C2) |
| 5. RNAscope™ Probe- Mm-Sst-C3 (ACD Cat No. 404631-C3) |
| 6. RNAscope™ Probe- Mm-Pvalb-C3 (ACD Cat No. 421931-C3) |
| 7. RNAscope Probe- Mm-Olig2-C3 (ACD Cat No. 447091-C3) |
| 8. RNAscope™ Probe-Mm-Calb1, (ACD Cat No. 428431) |
| 9. RNAscope™ Probe-Mm-Gad1-C3 , (ACD Cat No. 400951-C3) |
| 10. RNAscope™ Probe-Mm-Crym , (ACD Cat No. 466131) |
| 11. RNAscope™ Probe- Rhodopsin-ChR2-C3, (ACD Cat No. 508131-C3) |
| 12. RNAscope™ Probe-Mm-Dcn-C3, (ACD Cat No. 413281-C3) |
| 13. RNAscope™ Probe- Mm-Chat, (ACD Cat No. 408731) |
| 14. RNAscope™ Probe-Mm-Ache1-C3, (ACD Cat No. 556411-C3) |
| 15. RNAscope™ Probe-Mm-Fibcd1-C3, (ACD Cat No. 524021-C3) |
| 16. RNAscope™ Probe- Mm-Th , (ACD Cat No. 31762) |

B
| Immunofluorescence Reagents: |
| --- |
| 1. Cyclic AMP Primary Antibody (R&D Systems Cat. No. MAB2146) |
| 2. Recombinant Anti-PKA alpha/beta/gamma (catalytic subunit) (phosphoT197) antibody [EP2606Y] (Abcam Cat. No ab75991) |
| 3. M.O.M.® (Mouse on Mouse) Immunodetection Kit, Fluorescein ( Vector Laboratories Cat. No FMK-2201) |
| 4. Donkey Anti-Rabbit IgG (H+L), Highly Cross-Adsorbed CF568 (Biotium Cat. No #20098-1mg). |
| 5. Donkey Anti-Mouse IgG (H+L), Highly Cross-Adsorbed CF660R (Biotium Cat. No. #20388-1mg) . |
| 6. Normal Donkey Serum (Jackson ImmunoResearch Laboratories Cat. No 017-000-121). |
| 7. DAPI (4',6-Diamidino-2-Phenylindole, Dihydrochloride) (ThermoFisher Scientific Cat. No D1306) |

### Spiny Projection Neurons Highly Express *Gnal*

We first quantified the expression pattern of *Gnal* in the most abundant neuron type of striatum, SPNs, which abundantly express *Gnal* (Fig. 2). Expression in direct pathway SPNs, denoted by D_1_ dopamine receptor (*Drd1*) positive cell staining (Neve et al., 2004), averaged at 62.14 *Gnal* puncta per cell (Fig. 2B/C). Indirect pathway SPNs, labeled by D_2_ dopamine receptor (Drd2) positive cell staining (Neve et al., 2004), showed a similar *Gnal* expression pattern with an average of 65.7 *Gnal* puncta per cell (Fig. 2A/C). Expression of *Gnal* did not significantly differ between SPN classes (Fig 2C). During our characterization of the striatum, we also noted *Gnal* expression in the corpus callosum. In wake of this observation, we probed for oligodendrocyte transcription factor (*Olig2*) and confirmed oligodendrocytes of the corpus callosum lowly express *Gnal* transcripts (Sup. Fig. 3).

**Figure 2:**
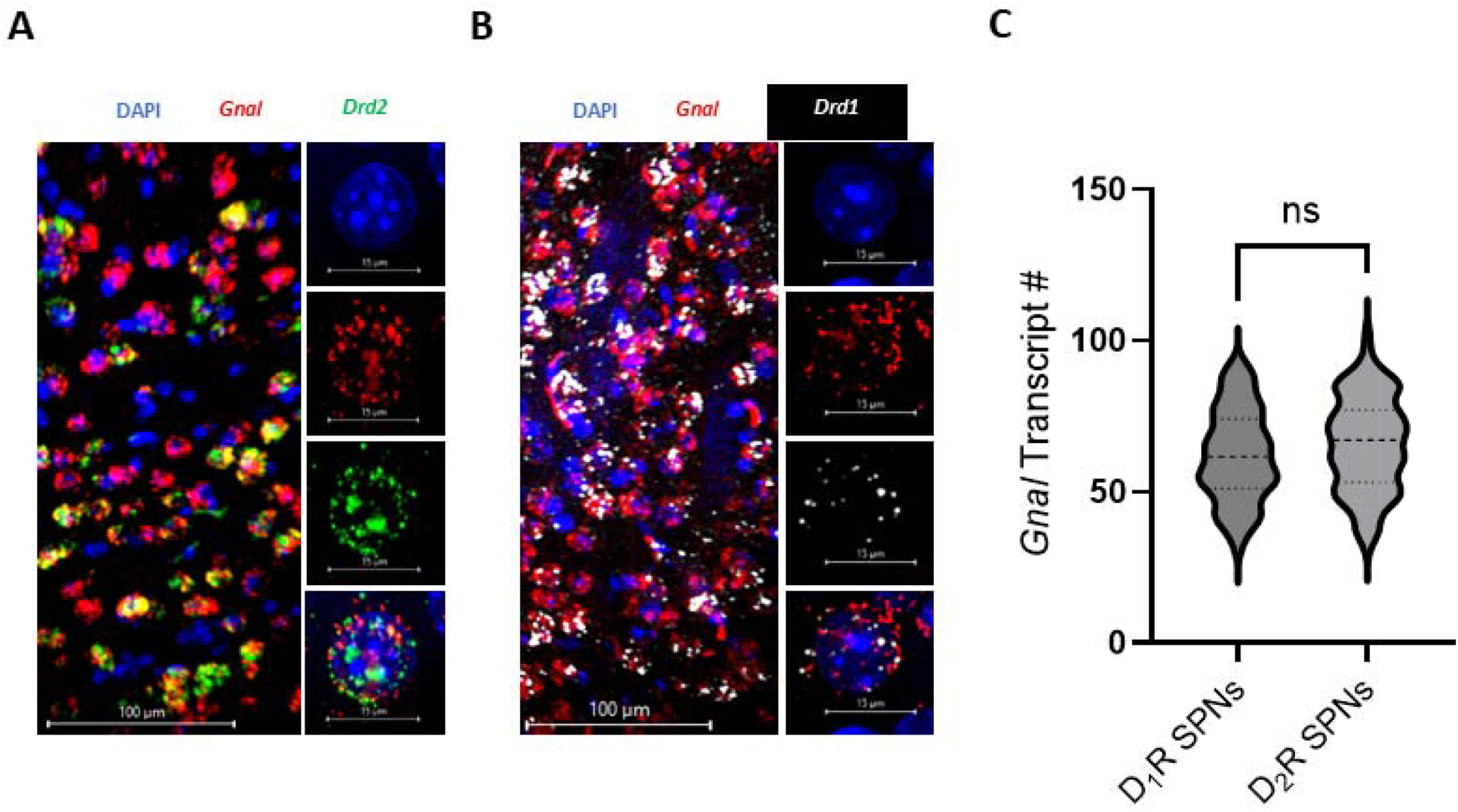
Spiny Projection Neurons Highly Express *Gnal*. A-B) Micrographs of *Gnal* (red) expression in spiny projection neurons (SPNs). The left most image depicts representative expression patterns of *Gnal* and biomarkers at 10x magnification. The right column of images shows signal for each channel of a representative cell at 60x magnification, merged at bottom. Nuclei are labeled by DAPI (blue). A) Micrographs depicting *Gnal* expression in *Drd2* (green) SPNs. B) Micrographs depicting *Gnal* expression in *Drd1* (white) SPNs. C) Single cell quantification of *Gnal* transcript expression. D_1_R SPNs average at 62.14 transcripts per cell (95% CI 59.14-65.14), and D_2_R SPNs average at 65.7 transcripts per cell (95% CI 62.61-68.79). Expression differences were not statistically significant; p>0.05, Unpaired-T test, t=1.639. N= 3 mice, n= 100 cells of each classification.

### Striatal Interneurons Differ in *G***α**-Encoding RNA Expression

Next, we characterized *Gnal* expression between interneuron populations in the striatum. *Gnal* transcription is enriched in cholinergic interneurons, with an average of 72 puncta, as seen by co-staining for murine Choline Acetyltransferase (*Chat*) and Acetylcholinesterase (*Ache1*) (Fig. 3A/D). In representative cells, *Gnas* transcript counts averaged 42.3 transcripts per cell (Fig. 4B), indicating cholinergic interneurons likely utilize Gα_olf_ as the major stimulatory G-protein alpha subunit.

**Figure 3:**
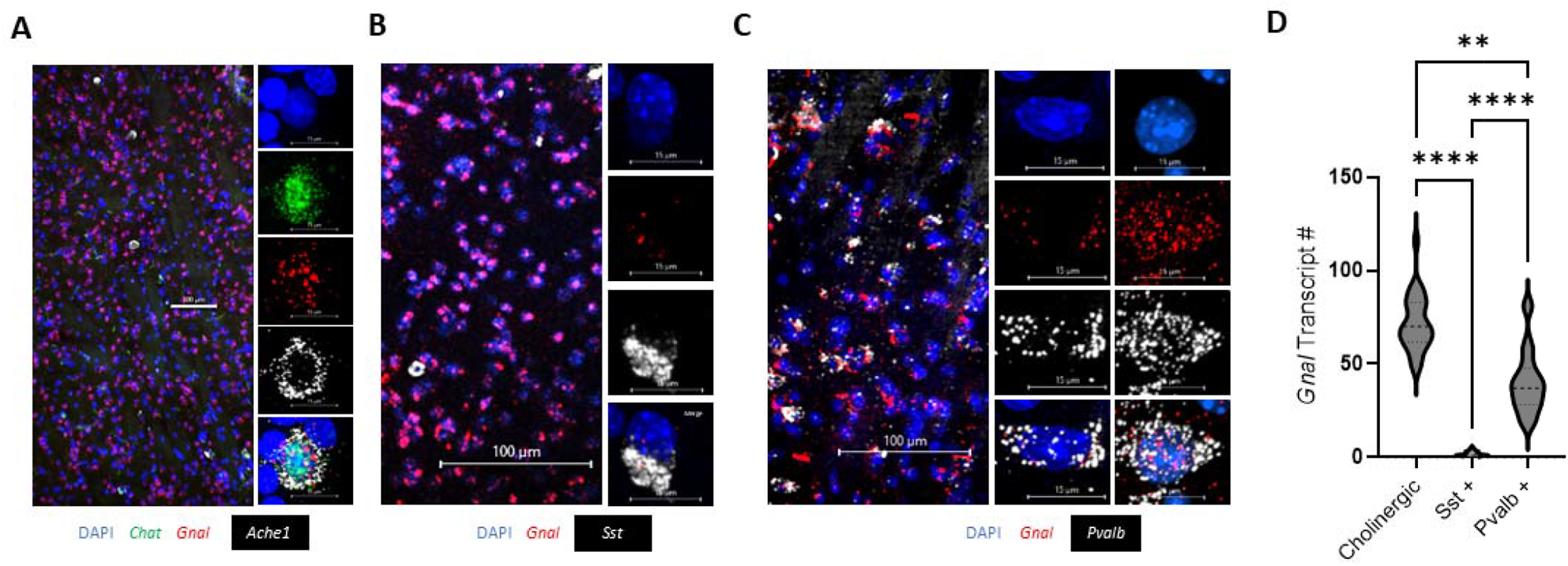
Striatal Interneurons Differ in *Gnal* Expression. A-C) Micrographs depicting *Gnal* (red) expression in interneuron populations. Left most image is 10x magnification, other columns of images depict representative cells at 60x magnification. A) Cholinergic interneurons, defined as *Chat* (green) and *Ache1* (white) positive cells, express *Gnal* transcripts. B) Somatostatin *(Sst*) (white) positive interneurons display low to no *Gnal* expression. C) Parvalbumin (*Pvalb*) (white) positive interneurons varied in *Gnal* expression. Both low expressing (middle column) and high expressing populations (right column) were observed. D) Quantification of RNAscope in situ hybridization. Cholinergic interneurons averaged at 72 transcripts per cell (95% CI 65.72-78.28), *Sst* positive interneurons 1.72 transcripts per cell (95% CI 1.55-2.285), and *Pvalb* positive interneurons 40.2 transcripts per cell (95% CI 33.26-47.14). Differences in *Gnal* expression between striatal interneurons are significant; Kruskal-Wallis statistic (H) = 61.31, P <0.0001. N= 3 mice, n = 25 of each cell type.

**Figure 4:**
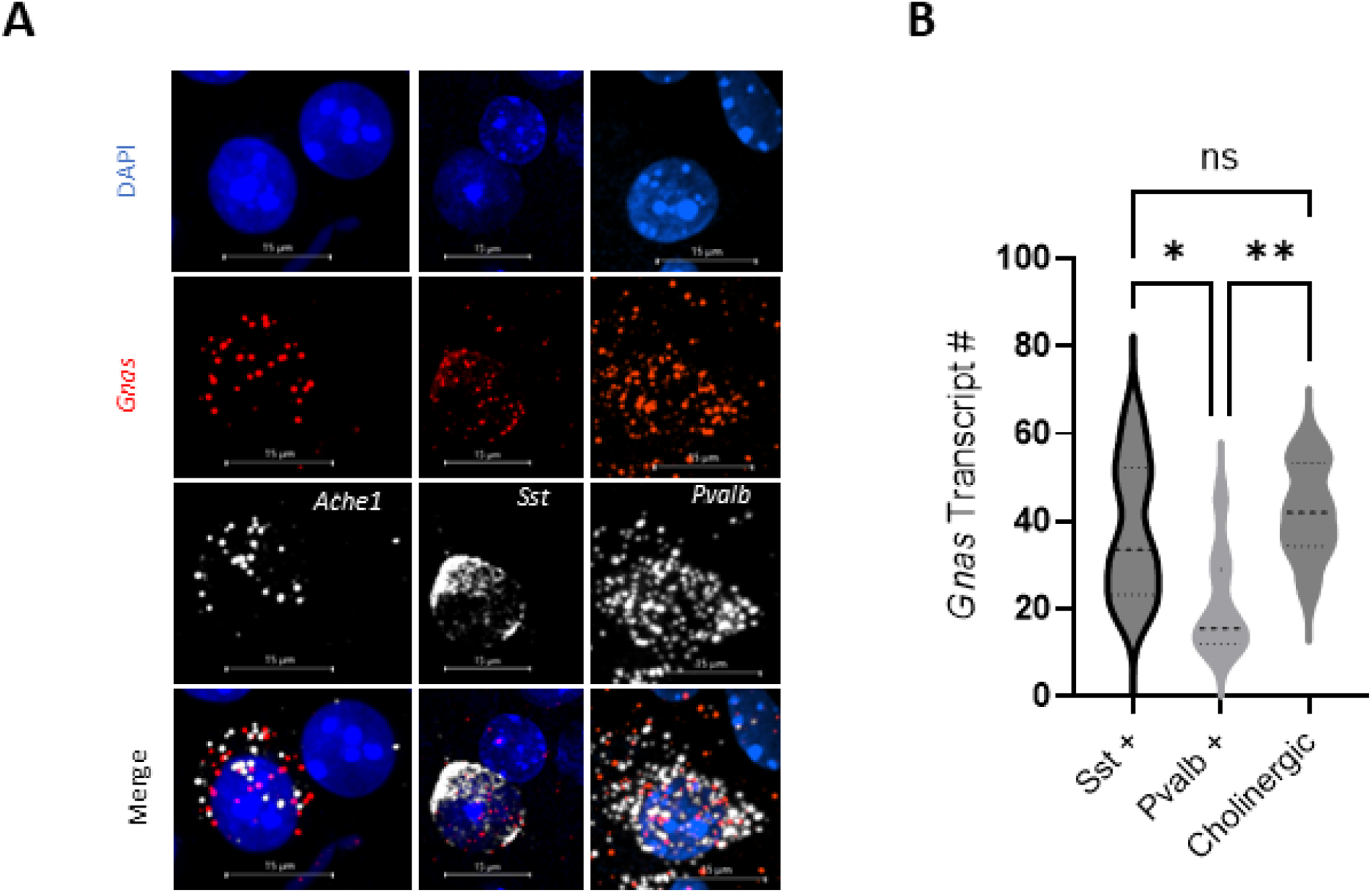
Striatal Interneurons Differ in *Gnas* Expression. A) Representative 60x micrographs of *Gnas* (red) expression. The left most column shows cholinergic interneurons, denoted by *Ache1* (white); middle column displays Somatostatin (*Sst) (*white) positive interneurons, and the right column displays *Pvalb* (white) interneuron populations. B) Single cell quantification of *Gnas* transcripts in three striatal interneuron populations. *(Sst*) positive interneurons displayed an average of 37.9 transcripts per cell (95% CI 26.68-49.12), *Pvalb* interneurons had a mean of 20.4 transcripts per cell (95% CI 12.41-28.39), and cholinergic interneurons 42.3 transcripts (95% CI 34.76-49.84). *Gnas* expression was statistically lower in *Pvalb* interneuron populations; Kruskal-Wallis statistic (H) = 11.90, P= 0.0026. N= 3 mice, n = 10 of each cell type.

It was initially noted that some striatal cells do not express *Gnal* transcripts. We hypothesized that the small subset of *Gnal* negative cells were either Somatostatin (Sst) or Parvalbumin (Pvalb) interneuron populations of the striatum. Upon co-staining for *Gnal* within *Sst* interneurons, we observed low to no *Gnal* transcripts per cell (Fig. 3B/D), with an average of 1.72 *Gnal* puncta per cell. To further define Gα signaling in this population, we co-probed for *Gnas* in representative cells and found moderate *Gnas* expression (Fig 4A). Quantification yielded an average of 37.9 *Gnas* transcripts per cell. This data suggests *Sst* positive striatal interneurons do not rely on Gα_olf_ as the major stimulatory heterotrimeric G-protein alpha subunit.

*Pvalb* positive interneurons were found to have largely variable *Gnal* expression (Fig. 3C), with an average transcript count was 40.2 transcripts per cell (Fig. 3D). Variability in this data can likely be attributed to multiple distinct interneuron populations expressing *Pvalb* (Muñoz-Manchado et al., 2018. Of the striatal interneurons sampled (Fig 4A, right column), we found comparatively lower *Gnas* transcript counts, averaging 20.4 transcripts per cell (Fig. 4B). Further characterization of Gα expression between subclasses of *Pvalb* interneurons may provide a more detailed understanding of stimulatory G-protein signaling between these interneurons.

### Layer 2/3 Cortical Neurons Express More *Gnal* Than Deep Cortical Layers

The first extrastriatal tissue we examined for *Gnal* expression was the cortex. *Gnal* expression was found to span all cortical layers but with enrichment in Layer 2/3 (Fig. 5A), as seen co staining for upper cortical marker Calbindin 1 (Calb1)(Zeng et al., 2012). Layer 2/3 neurons display an average of 47.22 transcripts per cell (Fig 5C). Deep cortical layers, determined by co-staining for Mu-crystallin (CryM)(Zeng et al., 2012) (Fig 5B) contain significantly less *Gnal* transcripts, with an average of 25.19 transcripts on a per cell basis (Fig. 5C). Cortical GABAergic cells were identified by Glutamate Decarboxylase 1 (*Gad1*) expression and populate all cortex layers (Sup. Fig. 4A). Cells that were *Gad1* positive were excluded from quantification of layer 2/3 and deeper cortical layer *Gnal* expression. The expression of *Gnal* transcripts within GABAergic cells the cortex was widely variable (Sup. Fig. 4B), even within cortical layers, likely due to the diverse population of GABAergic cells in the cortex (Gonchar et al., 2008).

**Figure 5:**
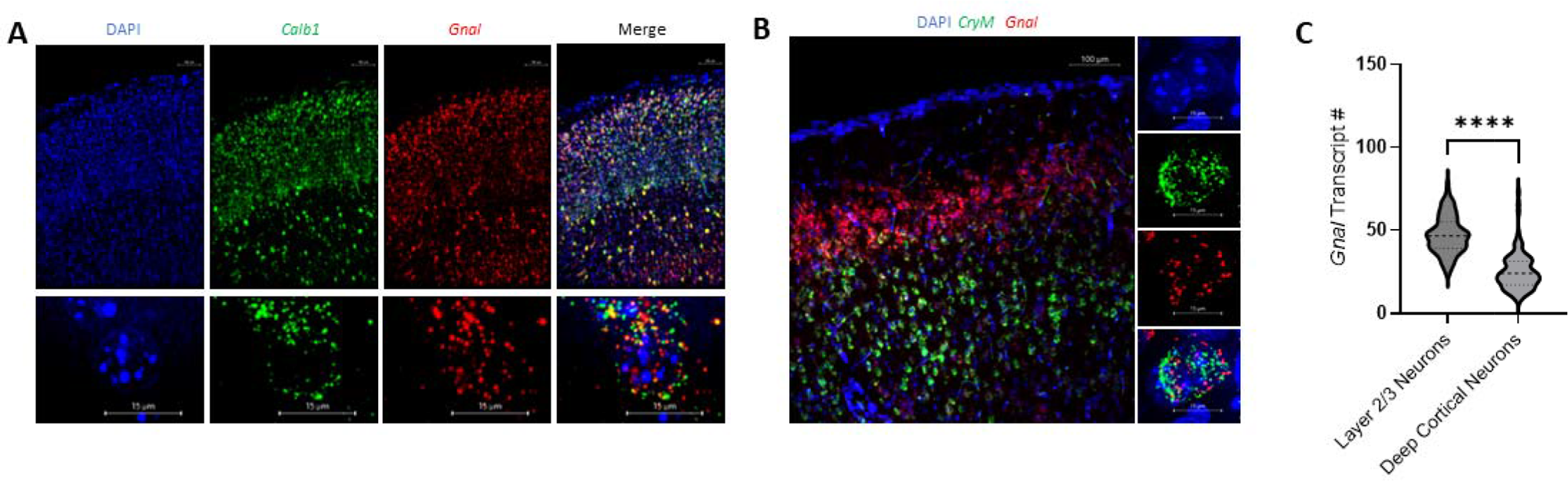
Layer 2/3 Cortical Neurons Moderately Express *Gnal*. A-B) Micrographs of *Gnal* RNA expression (red) in the cortex. A) The top row contains 10x images of mouse cortical layers and the bottom row depicts a representative cell at 60x magnification. Layer 2/3 neurons are denoted by *Calb1* (green) positivity and express *Gnal* transcripts (bottom right). B) 10x magnification of mouse cortex (left), with deep cortical layers marked by *CryM* (green) positivity. Representative CryM positive cells at 60x magnification (right column) show *Gnal* expression. **C)** Quantitative analysis of single cell *Gnal* transcript expression in cortical neuron populations. Layer 2/3 neurons display moderate *Gnal* expression, with an average of 47.22 transcripts per cell (95% CI 44.98-49.96). Deep cortical layers also express *Gnal* at statistically lower per-cell transcript counts, averaging 25.19 transcripts per cell (95% CI 22.91-27.48). p< <0.0001, Unpaired-T test, t= 13.65. N= 3 mice, n = 100 Layer 2/3 neurons, 98 deep cortical neurons.

### Dopaminergic Cells of the Midbrain Moderately Express *Gnal*

Next, we characterized *Gnal* expression in the substantia nigra. Dopaminergic cells of the substantia nigra pars compacta (SNc), marked by positive staining for tyrosine hydroxylase (Lammel et al., 2015) (Fig. 6A), were found to express an average of 47.43 *Gnal* puncta per dopaminergic neuron (Fig. 6B). The substantia nigra pars reticulum (SNr), histologically identified with *Gad1* staining directly below the SNc (Fig. 6A), displays lower expression levels of *Gnal* expression, with an average of 24.20 puncta per cell (Fig. 6B). As the SNc and SNr overlap depending on stereotaxic coordinates, SNc cells were defined as *Th* positive but *Gad1* negative and visceral for the SNr. The ventral tangential area (VTA) also displays positive *Gnal* staining but was not quantified in this project (Fig. 6A).

**Figure 6:**
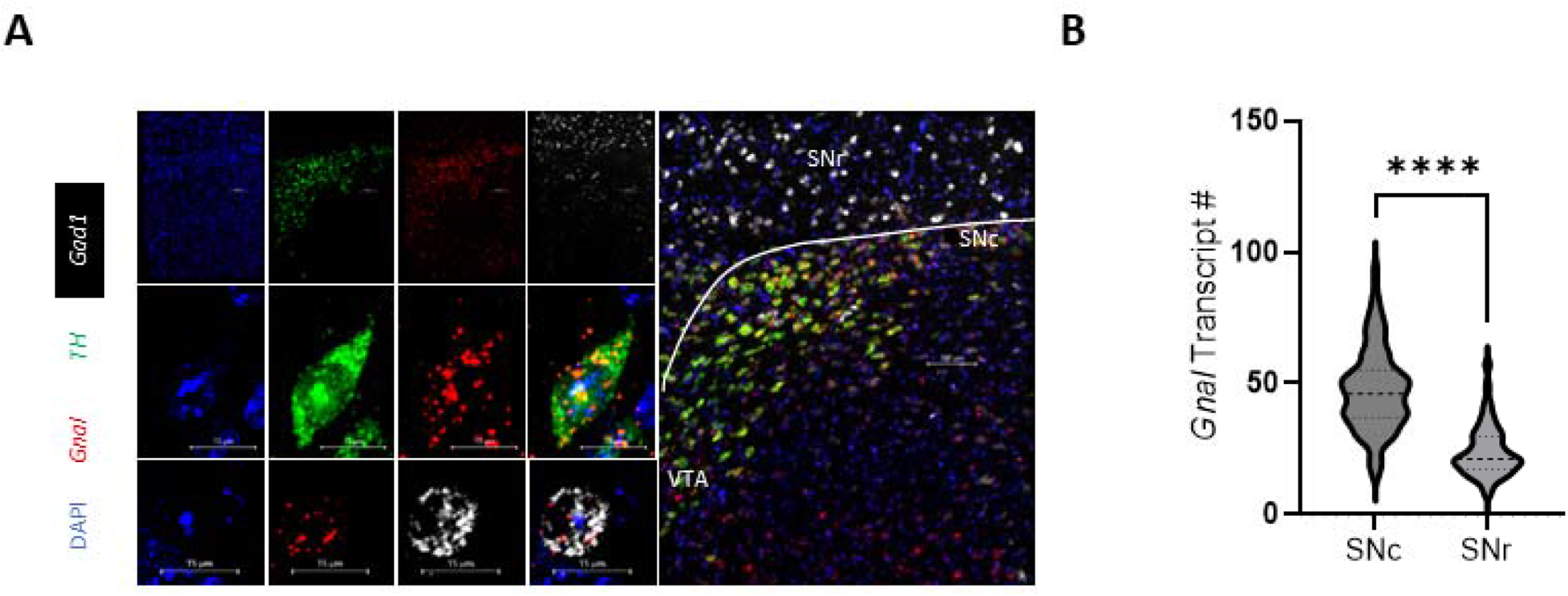
Dopaminergic Cells of the Midbrain Moderately Express *Gnal*. A) Micrographs depicting *Gnal* (red) expression in the midbrain. Top row are 10x magnification images, which are merged and enlarged in the right most image. The middle row and bottom row are 60x magnification images of representative *TH* (green) and *Gad1* (white) neurons. B) Dopaminergic cells of the substantia nigra pars compacta (SNc) displayed an average of 47.43 transcripts per cell (95% CI 44.31-50.56), and GABAergic cells of the substantia nigra pars reticulata (SNr) averaged at 24.20 transcripts per cell (95% CI 22.16-26.23). *Gnal* expression was statistically different between the two groups; p< 0.0001, Unpaired-T test, t= 12.17. N= 3 mice, n= 99 SNc neurons, 92 SNr neurons.

### Hippocampal Neurons Lowly Express *Gnal*

Our initial characterization showed relatively strong *Gnal* signal in hippocampal subfields CA1, CA2, and CA3 as well as the dentate gyrus (Fig.1A). To further characterize this expression, we utilized Fibrinogen C Domain-Containing Protein (*Fibcd1*), which is vastly enriched in hippocampal subfields and dentate gyrus (Cembrowski et al., 2016), providing staining patterns clear enough to enable its use as a biomarker of both structures (Fig. 7A). Due to the high cell density of the dentate gyrus, *Gnal* appears to be highly expressed but single cell expression average of 12.76 transcripts per cell (Fig. 7B). Hippocampal subfields CA1, CA2, CA3 average 12.84 *Gnal* transcripts per cell (Fig. 7B). No obvious differences in expression were noted between subfields and they were quantified as one entity.

**Figure 7:**
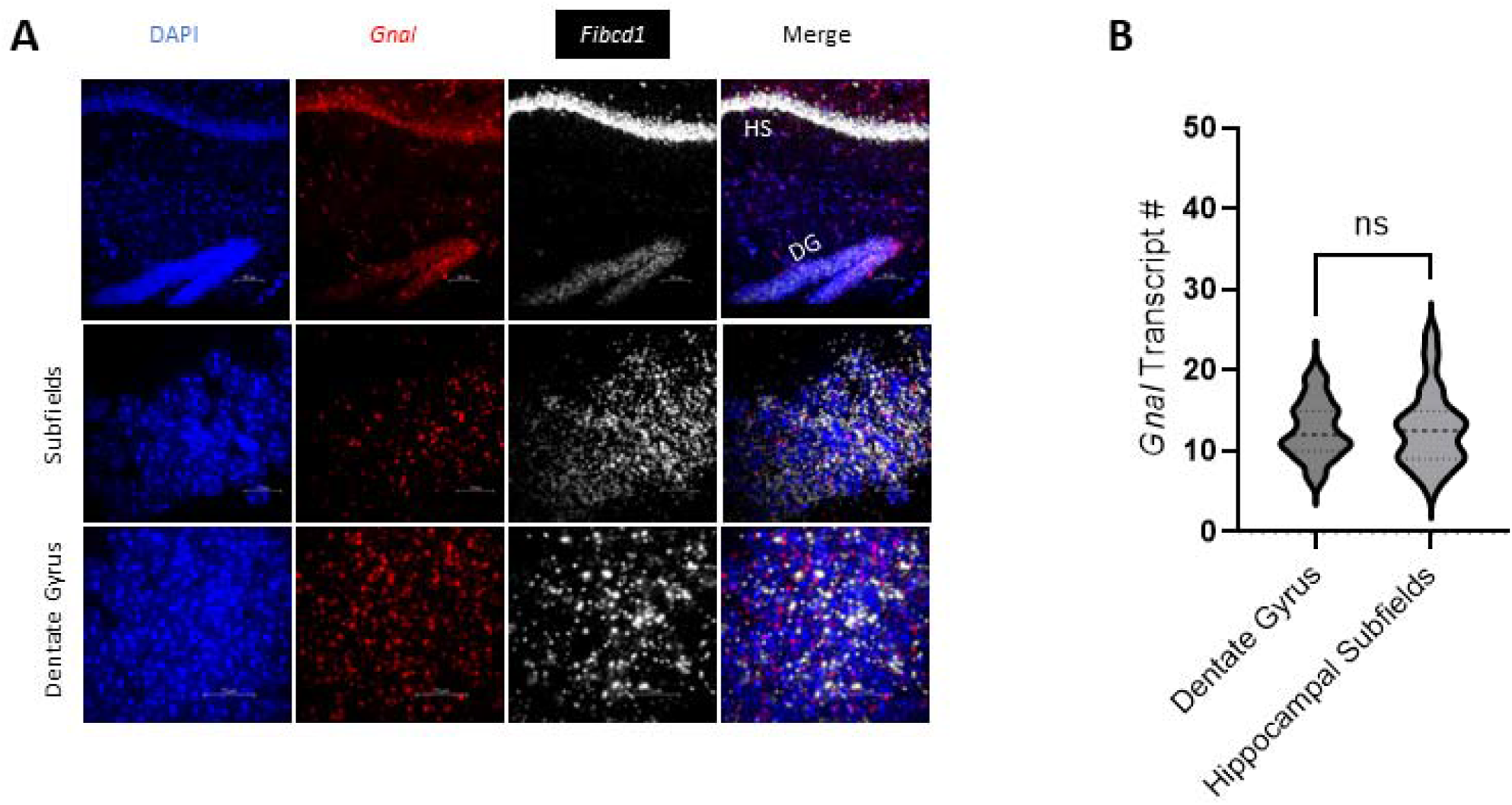
Hippocampal Neurons Lowly Express *Gnal*. A) Micrographs of *Gnal* expression (red) in the hippocampus at 10x magnification. *Fibcd1* (white) expression was used to identify hippocampal subfields and the dentate gyrus. B) Quantitative analysis of single cell *Gnal* transcript expression. The dentate gyrus (DG) averaged at 12.76 transcripts per cell (95% CI 12.04-13.47) and hippocampal subfields (HS) at 12.84 transcripts (95% CI 11.91-13.77). Expression differences were not statistically significant; p>0.05, Unpaired-T test, t=0.1433, N= 3 mice, n = 98 DG neurons, 100 HS neurons.

We also observed sparse amounts hippocampal cells outside of these structures that have moderate *Gnal* expression, and subsequently co-probed for Oriens-Lacunosum/Moleculare (OLM) interneurons. Representative OLM interneurons, identified by *Sst* positivity (Speigel & Hemmings Jr, 2022), were imaged and found to have low levels of *Gnal* expression (Sup. Fig. 5). Neighboring cells express substantially more *Gnal* (Sup Fig. 5 white arrow), and further studies are necessary to characterize expression between the large diversity of hippocampal interneurons (Booker & Vida, 2018).

### Purkinje Cells Highly Express *Gnal*

Lastly, we defined the expression of *Gnal* in the cerebellum. Cerebellar *Gnal* is largely restricted to the Purkinje cell layer of the cerebellum (Fig. 8A/C). Purkinje cells express the highest number of *Gnal* transcripts out of any cell analyzed in this study, averaging 119.3 transcripts per cell (Sup. Fig. 6). The dense granule cell layer of the cerebellum expresses single digit *Gnal* transcripts per cell, with an average of 2.876 transcripts (Fig. 8A/C). The molecular layer of the cerebellum displays sparse *Gnal* positivity but was not included in the characterization. Co-probing representative Purkinje cells for both *Gnal* and *Gnas* expression unveiled a large difference in expression, with average *Gnas* expression of 36.10 transcripts per cell (Fig. 8D). Gnal transcript numbers were consistent with previous analysis.

**Figure 8:**
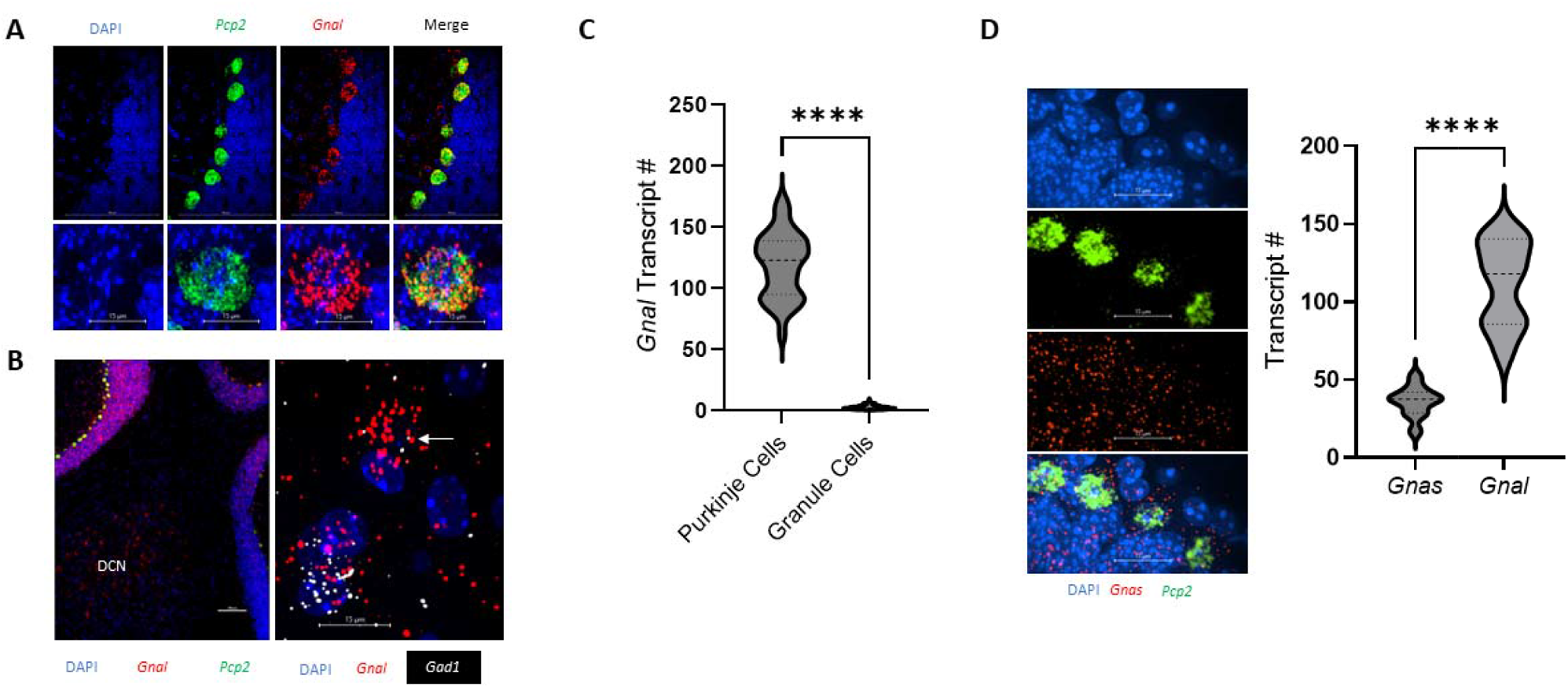
Purkinje Cells Highly Express *Gnal*. A-B) Micrographs of *Gnal* (red) RNA expression within the cerebellum. A) Purkinje cells, identified by *Pcp2* (green) expression, show *Gnal* expression. The top row is 10x magnification and bottom row is a representative Purkinje cell at 60x magnification. B) Micrograph depicting *Gnal* expression localized to deep cerebellar nuclei (DCN) at 10x magnification (left) and 60x magnification (right). *Gad1* (white) positive DCN cells express less *Gnal* transcripts compared to neighboring unknown cell (white arrow). C) Quantitative analysis of single cell *Gnal* transcript expression in the cerebellum showed an enrichment of transcripts in Purkinje cells with an average of 119.3 transcripts per cell (95% CI 114-124.6). The granule layer presents lower *Gnal* expression, with a mean of 2.876 transcripts per cell (95% CI 2.538-3.215). *Gnal* expression was statistically significant between cell types, p< 0.0001, Unpaired-T test, t=43.03. N= 3 mice, n = 100 Purkinje cells, 97 granule cells. D) Micrograph depicting *Gnas* expression patterns within the cerebellum at 10x magnification (left) and transcript quantification in representative Purkinje cells (right). Average *Gnas* expression observed was significantly lower (p< 0.0001, Unpaired-T test, t= 8.298) at 36.10 transcripts per cell (95% CI 29.17-43.03). N= 3 mice, n= 10 Purkinje cells.

Deep cerebellar nuclei (DCN) also express *Gnal* at varying transcript numbers (Fig. 8B). We co-probed for *Gad1* and found GABAergic cells of the DCN only lowly expressed *Gnal* (Fig 8B, right image). Other higher expressing cell types were observed (Fig. 8B, white arrow). Further experimentation will be necessary to elucidate the differential *Gnal* expression between DCN cell types.

### Loss of *Gnal* Causes Molecular Deficits in Purkinje Cell Layer

Cerebellar *Gnal* expression data suggested that Gα_olf_ may play a major role in cAMP production in Purkinje cells. To test this idea, we probed both wild type and Purkinje cell conditional *Gnal* knockout tissue with an antibody specific to cAMP. Subsequent imaging of cerebellar Purkinje cells in both tissue types revealed loss of *Gnal* leads to significant decreases in intracellular cAMP fluorescent signal (Fig. 9B).

**Figure 9:**
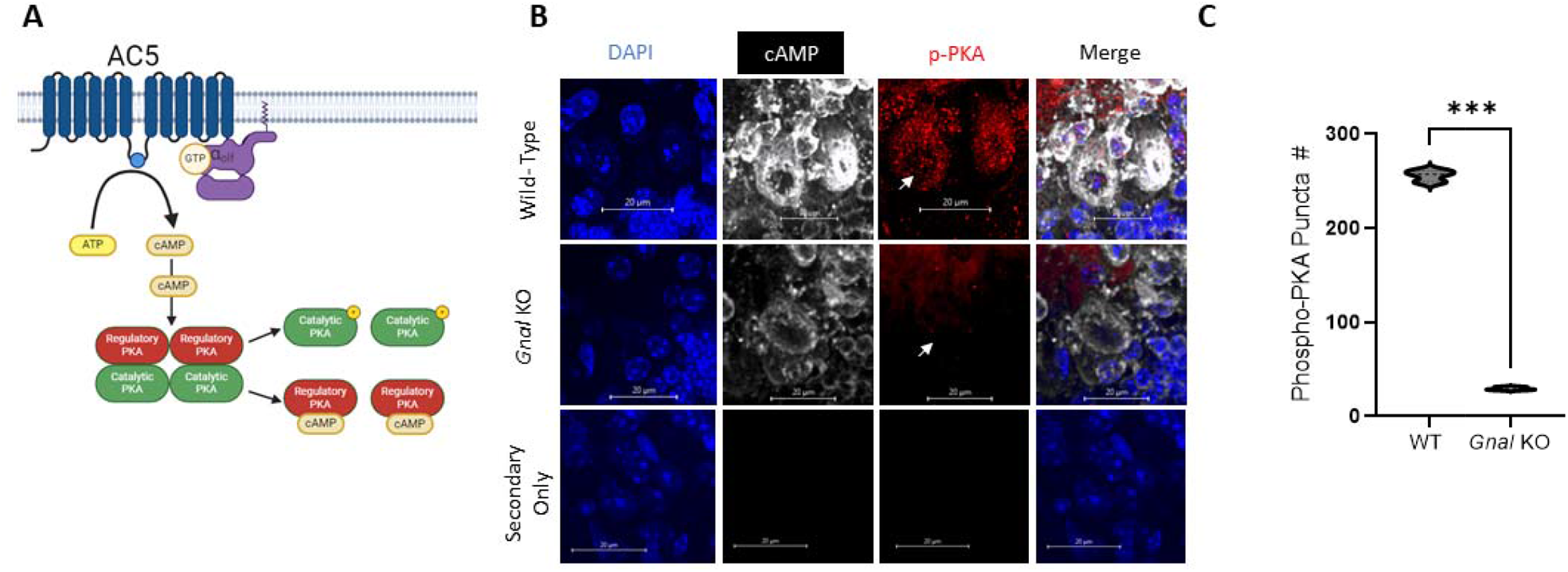
Loss of *Gnal* results in Purkinje Cell Molecular Dysfunction. A) Schematic showing Gα_olf_ stimulation of adenylyl cyclase 5 (AC5) mediated cAMP production and downstream cAMP activation of protein kinase A. ATP is first converted into cAMP by AC5. Next cAMP binds the regulatory subunits of PKA allowing the catalytic subunits to dissociate and become fully active via phosphorylation at threonine 197 (depicted as yellow P in diagram). **B)** Micrograph of representative Purkinje cells at 60x magnification. Wild-type cerebellar Purkinje cells (top row) display strong signal for intracellular cAMP (white) and activated PKA (red). *Gnal*-knockout cerebellar Purkinje cells (middle row) display comparatively less cAMP and activated phospho-PKA signal (p-PKA). **C)** Quantification of active PKA puncta between wild-type (WT) and *Gnal* KO tissue. WT tissue averaged at 255.7 puncta (95% CI 240.5-270.8) per Purkinje cell and *Gnal* KO tissue at 29.7 puncta per cell (95% CI 25.54-33.13). Differences in puncta quantification were significant; p = 0.0001, Welch’s T test, t=62.24. N= WT mice, 2 Purkinje cell *Gnal* KO mice, n= 3 Purkinje cells.

Intracellular cAMP is known to associate with protein kinase A (PKA), which is a tetramer composed of two regulatory and two catalytic subunits (Turnham & Scott, 2016). Upon binding of cAMP to the PKA regulatory subunits, PKA catalytic domains dissociate and are phosphorylated at Threonine 197 to become fully active (Cheng et al., 1998) (Fig. 9A). To determine the consequences of Gα_olf_loss on protein activation downstream of cAMP production, we next performed antibody labeling for the activated catalytic subunit of PKA. Signal for active PKA catalytic subunits shows substantial differences between wild-type controls and Purkinje cell *Gnal* knockout (Fig 9B). Quantification of representative cells revealed wild type tissue averaged at 255.7 puncta per cell and *Gnal* KO tissue at 29.7 puncta per cell (Fig 9C).

## Conclusions

Gα*_olf_*expression and signaling has been best described within SPNs of the striatum, and many studies of *GNAL* linked dystonia have focused on the striatum. Our results characterize *Gnal* expression in neuronal cell populations throughout the brain, and we found some degree of *Gnal* expression in most brain regions examined. This raises fundamental questions surrounding Gα_olf_ signaling in different parts of the brain, and consequently how mutations in *Gnal* lead to dystonia.

Though it has previously been noted that not all striatal cells express *Gnal* (Herve et al., 2001), our results show new insights into the expression pattern of *Gnal* in striatal neurons. In SPNs, both direct and indirect pathway SPNs had high numbers of *Gnal* transcripts, and there were no statistically significant differences in the expression of *Gnal* between SPN subtypes. This reaffirms that Gα_olf_ may play a similar important role in stimulatory Gα signaling in both SPN classes. Additionally, we observed that cholinergic interneurons express higher number of *Gnal* transcripts than SPNs. Probing for *Gnas* in striatal cholinergic interneurons also showed significantly lower transcript number compared to *Gnal*. This shows that not only is Gα_olf_ the predominant stimulatory Gα isoform in cholinergic cells, but also has the highest transcript number in the striatum. This points to a potentially unique role for Gα_olf_ in modulating cholinergic interneuron physiology, and reaffirm that striatal cholinergic interneurons may play a critical role in the pathophysiology of dystonia. Furthermore, we report for the first time that *Sst* expressing striatal interneurons express low to no *Gnal* transcripts, and that *Pvalb* expressing interneuron populations show a variable *Gnal* expression pattern that likely differs based on subclassification of *Pvalb+* interneurons. (Muñoz-Manchado et al., 2018). Taken together, our data point to Gα_olf_ signaling being present to most neuron types in the striatum, and that mutations in *GNAL* may drive dystonia phenotypes through multiple mechanisms both within the microcircuitry of the striatum and through changes within projection neurons from the striatum themselves.

Cerebellar Purkinje cells were found to express the highest number of *Gnal* transcripts per cell, and *Gnas* transcripts were substantially lower compared to *Gnal*, suggesting Gα_olf_ is the predominant activator of adenylyl cyclase mediated cAMP production in these cells. This was confirmed with our use of a novel mouse model to knockout *Gnal* in Purkinje Cells, and knockout animals had significantly lower intracellular cAMP levels and phosphorylated PKA than controls. This shows that Gα_olf_ signal transmission is important to Purkinje cell cAMP-dependent pathways. Downstream of Purkinje cells are cells of the deep cerebellar nuclei. We show that there is differential expression of *Gnal* in this nucleus with *Gad1+* cells showing no to low expression, and *Gad1-* negative cells showing robust expression. These cells are likely glutamatergic projection neurons from this nucleus which have recently been shown to regulate dopamine release from dopaminergic cells of the SNc (Yoshida et al., 2022). These data show that Gα_olf_ function in cerebellar circuits could represent extra-striatal pathways to influence dystonia motor phenotypes caused by *GNAL-*linked mutations, and this is supported by recent evidence in *Gnal* haploinsufficient rodent models(Aïssa et al., 2022).

Outside of the striatum and cerebellum, we have observed new insights into *Gnal* expression in several neuron types that either are part of the basal ganglia processing loops or are known regulators of basal ganglia processing. In the cortex, we found modest *Gnal* expression in Layer 2/3 pyramidal neurons and lower, sparse expression in some deep cortical layer pyramidal neurons. Both cortical layers have variable expression of *Gnal* in *Gad1+* interneurons. Like the striatum, this suggests Gα_olf_ signaling may directly influence cortical signaling through multiple mechanisms including through microcircuit based mechanisms, cortical processing of inputs, and altering projection neurons themselves. We did not observe differences in *Gnal* expression between cortices, suggesting Gα_olf_may be broadly influence cortex functions There have been no studies that directly examine Gα_olf_signaling either in the context of normal physiology or in *GNAL* linked dystonia, and may suggested an unexplored mechanism of *GNAL* linked dystonia.

Furthermore, we have found that several other basal ganglia structures express *Gnal.* SNc dopaminergic neurons express high transcript numbers for *Gnal*, and suggests striatal projecting SNc dopaminergic neurons may be an additional mechanism that Gα_olf_can modulate motor control (Chambers et al., 2023). The main output nucleus of the rodent basal ganglia, substantia nigra pars reticulata also expresses *Gnal*. Together, our investigation of *Gnal* expression shows that there are multiple points in the basal ganglia that are influenced by Gα_olf_ signaling, there are multiple points that corrupted Gα_olf_ signaling could initiate dystonic phenotypes from, or that abnormal Gα_olf_ signaling induced by mutations in *GNAL* lead to dystonia through altered network activity from multiple basal ganglia nuclei.

Beyond expression of *Gnal/*Gα_olf_, our data raise questions about how Gα_olf_ signals throughout the brain. To our knowledge, the GPCRs to which Gα_olf_ couples is unknown outside the context of striatal SPNs. There is large diversity of GPCR expression between brain regions (Azam et al., 2020) and identifying the specific interactions will be crucial to understanding which signal Gα*_olf_* transduces in various cells. Gα*_olf_* is currently described in binding AC5, but the expression of adenylyl cyclase isoforms also varies between brain tissue (Devasani & Yao, 2022), making novel AC-Gα_olf_ interactions likely in extra-striatal tissue, also raising the possibility of Gα_olf_ interacting with non AC proteins that are not currently known. Moreover, the expression of Gβγ subtypes is widely variable between brain tissue (Tennakoon et al., 2021), and Gα_olf_is currently characterized in binding to Gβ_2_ and Gγ_7_ isoforms (Schwindinger et al., 2010). Our *Gnal* expression data and expression patterns of Gβ_2_ and Gγ_7_ are not identical, suggesting unique combinations of Gα_olf_ and Gβγ isoforms in some tissue.

All together our data has raised many exciting possibilities for how *GNAL/*Gα_olf_influences neuronal signaling and activity. The widespread expression of *Gnal* transcripts in murine brain tissue implicates Gα_olf_in a diverse set of signaling pathways across many neuronal cell types. Due to the ubiquitous expression of *Gnal* in murine tissue, we hypothesize that dystonia linked *GNAL* mutations result in pathogenic signaling in many neuroanatomical structures, some of which may contribute to motor symptoms. Further investigation with our novel conditional *Gnal* flx/flx mouse model reported here will help to provide mechanistic understanding of how dystonic Gα_olf_ signaling leads to motor symptoms.

## Experimental Procedures

### Tissue Processing

Three wild type C57BL/6 adult mice were anesthetized with isoflurane and subsequently transcardial perfused with ice cold PBS and 4% PFA. After perfusion, the intact brains were recovered and post-fixed in 4% PFA for another 24 hours at 4°C. Samples were then sectioned into 15 µm thick slices on Leica vt1200s vibratome in ice cold PBS. Each slice was mounted onto a Superfrost slide, dried, and placed in a slide box with desiccant for short term storage at −80 Celsius.

### RNAScope In Situ Hybridization

RNAscope was performed according to the manufacturer’s recommended workflow. Briefly, 15 µm mouse brain slices were thawed in PBS, baked at 60°C, dehydrated in ethanol, and treated with hydrogen peroxidase before boiling in target retrieval buffer. Slides were then briefly immersed in ethanol and a hydrophobic barrier was drawn around the sample. Incubation with RNA probes was performed at 40°C, and samples were subjected to three amplifier hybridization incubations before fluorescent signal was developed for each channel. DAPI counterstaining occurred directly before applying a permanent coverslip. Reagents are listed in Table 1A.

### Immunofluorescence

Slices from all three mice were subjected to free floating *Gnal* antibody labeling in the wells of a 24-well plate. Samples were first blocked and permeabilized in PBS with 5% normal donkey serum and 0.1% triton x-100 for 2 hours at room temperature. *Gnal* primary antibody was diluted 1:100 in PBS with 5% donkey serum before being applied to samples for overnight incubation. The next morning, samples were washed three times in PBS and incubated with secondary antibody at room temperature for two hours. Three more PBS washes were performed before counterstaining with DAPI for 5 minutes. Samples were moved to a glass bottom dish for spinning disc confocal imaging. Concurrent sets of samples were processed without primary antibody incubations to serve as secondary-only controls.

Cyclic-AMP antibody (R& D Systems) labeling was performed on cerebellar tissue as above with two modifications to reduce off target staining associated with the use of mouse primary antibodies on mouse tissue. First, tissue was incubated with Mouse-on-Mouse (M.O.M) Blocking reagent (Vector Laboratories) for 1 hour at room temperature after the traditional blocking step. Second, both the primary and secondary antibodies were diluted in M.O.M diluent (Vector Laboratories) instead of traditional antibody diluent. Reagents are listed in Table 1B.

### Knockout of *Gnal* in Purkinje Cell Layer of Mice

Creation of the novel *Gnal* floxed mouse was achieved through a partnership with Jackson Labs Model Generation Service. Briefly, CRISPR/Cas9 gRNA constructs were used to insert loxP sites downstream of Exon 3 and upstream of exon 4. Mice were backcrossed to be congenic on C57Bl/6J and are maintained via flx/flx to flx/flx breeding. Genotryping was done through Transnetyx, Inc. Genetic knockout of *Gnal* in the Purkinje cell layer of C57BL/6J mice was accomplished by crossing our *Gnal* flx/flx mice with a Purkinje cell specific Cre line (L7-Cre, Jackson Laboratory, Strain #: 004146). Mice were bred and maintained as L7-Cre +/− *Gnal* flx/flx. Cre was always kept on the female breeder. Mice were used for experiments at >3months of age.

### Imaging

In-situ hybridization images were acquired on a Nikon spinning disc confocal with a 10x NA 0.3 objective for qualitative analysis of RNA expression patterns. All images quantified were acquired using a 60x NA 1.4 oil objective with 0.3 um step size z-stack sectioning. The 405 laser was set to 2.4% to visual DAPI staining in all samples. The 561 laser was set to 6.9% to quantify *Gnal* puncta. A consistent exposure time of 500 ms was used for image acquisition. Using these settings, less than 1 nonspecific puncta dot was detected per 10 cells using the negative 3-plex probe (Sup. Fig. 2A) and clear signal for all 3-plex positive control channels were observable (Sup. Fig. 2A). We noted the tyramide signal amplification (TSA) dyes had affinity to bind blood vessels in murine tissue, as seen in both positive and negative control samples (Sup. Fig. 2A). Off target blood vessel staining did not interfere with quantitative analysis.

Immunofluorescence images were also captured Nikon spinning disc confocal with a 10x NA 0.3 objective. Laser power and exposure time was based off secondary only controls and maintained consistent across all samples comparatively imaged. For low magnification images of *Gnal* tissue distribution in Figure 1 and Supplemental Figure 1, we used a Nikon AZ100 Multizoom Microscope with an RFP filter.

### Post Processing and Quantification

IMARIS software (Oxford Instruments) was used to process and quantify all images. First, a 3D reconstruction of each z-stack image was created in “3D view” tab of IMARIS. The number of *Gnal* RNA puncta was counted within cell of interest using the “spot analysis” function in IMARIS. Analysis settings were chosen from representative *Gnal* puncta measurements, and the processing settings were maintained consistent across all samples. The “surfaces” function was used to help define cell boundaries for the various biomarkers used.

## Acknowledgements

We would like to thank the laboratory of Dr. Jonathan Bird for allowing us to complete the large portion of imaging for this project on their Nikon Spinning Disc Confocal, and the laboratory of Dr. Nikhil Urs for allowing us to use their Nikon AZ100 fluorescent imager. Additionally, we would like to thank Tyler’s Hope for Dystonia and the National Institutes of Health and National Institute of Neurological Disorders and Stroke R00110878 to MSM for funding this work.

## Data Availability

All raw data, analyses, and documentation will be available upon request to the corresponding author.

## Ethics Statement

All animal studies were approved by UF IACUC.

