## Supplemental Figures for "*G*α*_olf_* Regulates Biochemical Signaling in Neurons Associated with Movement Control and Initiation"

**Supplemental Figure 1: Antibody Labeling Confirms the Translation of *Gnal* transcripts into G$\alpha_{olf}$ protein. A)** Micrograph depicting G$\alpha_{olf}$ expression (red) in a representative coronal slice of C57BL/6J brain tissue. **B)** Enlarged expression patterns in regions of interest for this study.

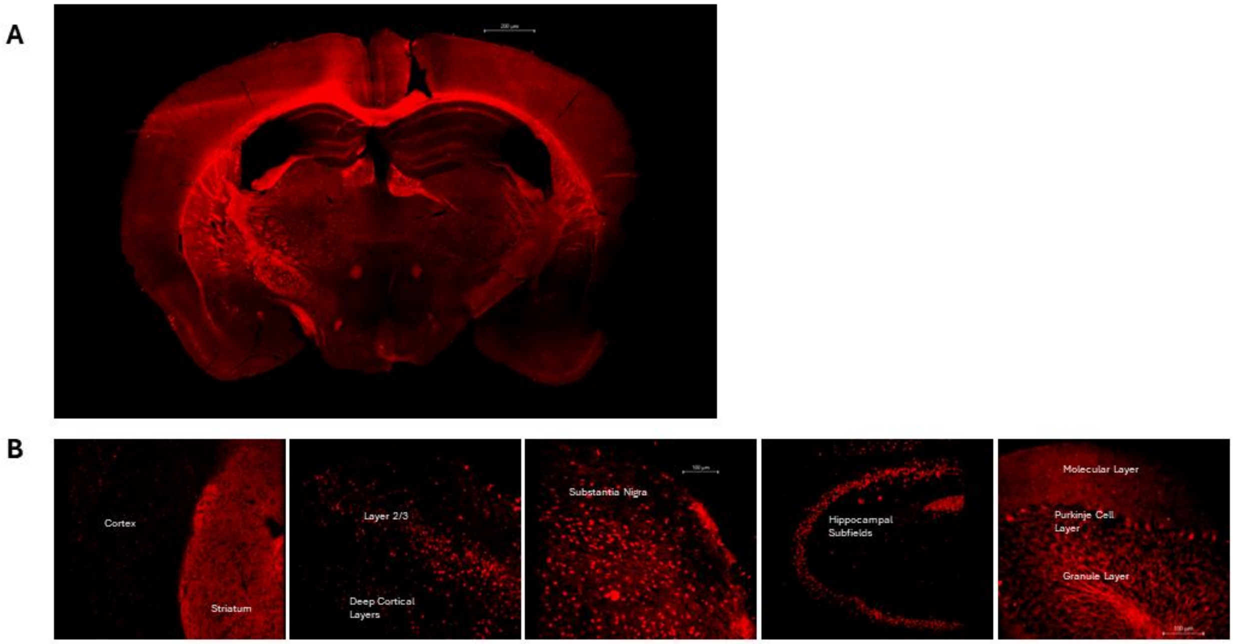

**Supplemental Figure 2: RNAScope 3-Plex Controls Define Acquisition Parameters. A)** Micrographs of RNAScope 3-plex positive control (top row) and 3-plex negative control (bottom row) at 10x magnification. The positive control contains probes for RNA polymerase II subunit A (*Polr2a*) in channel 1 (C1), peptidylprolyl isomerase B (*Ppib*) in channel 2 (C2), and ubiquitin C (*Ubc*) in channel 3 (C3). As seen in C1 of the negative control image, TSA dye binds to blood vessels in murine brain tissue, largely in the green channel. Off target staining of blood vessels did not interfere with quantitation of transcripts.

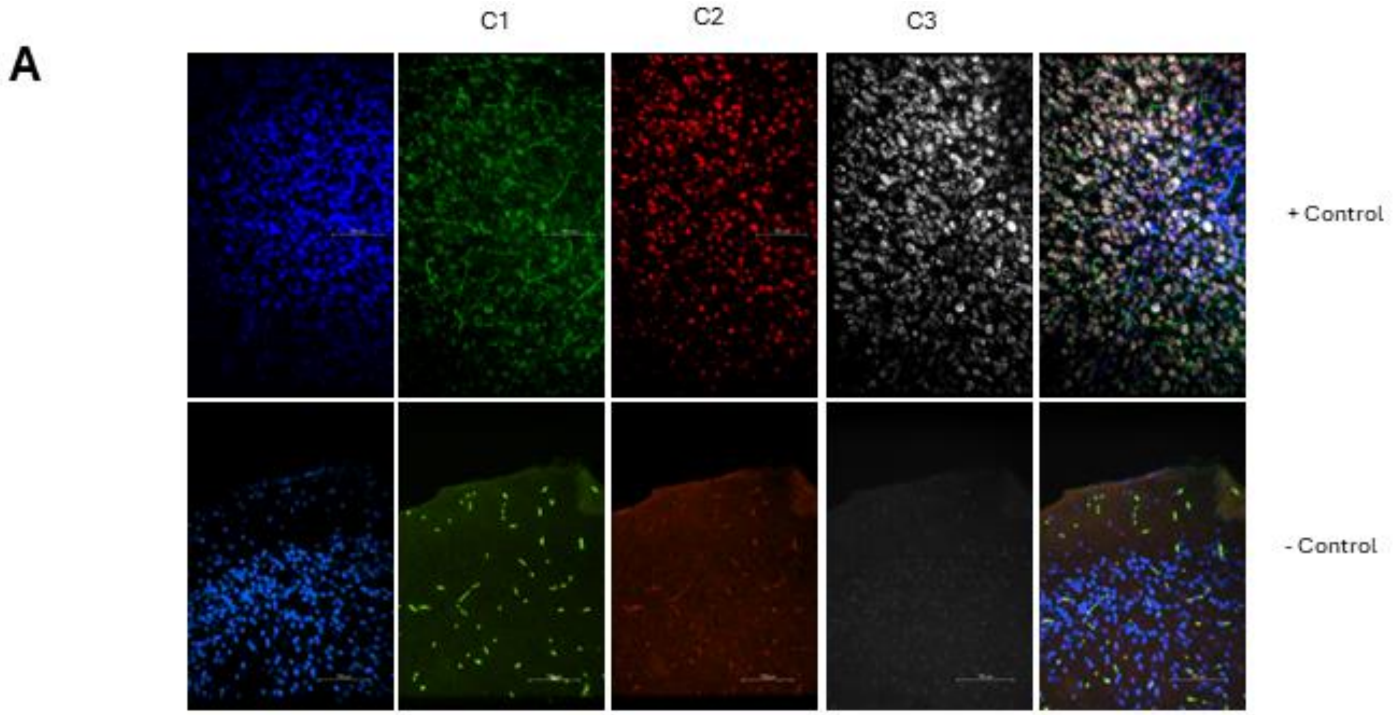

**Supplemental Figure 3: Corpus Callosum Oligodendrocytes Lowly Express Gnal. A)** Micrographs of *Gnal* expression in *Olig2* positive cells of the corpus callosum at 10x (left) and 60x (right column) magnification.

**A**

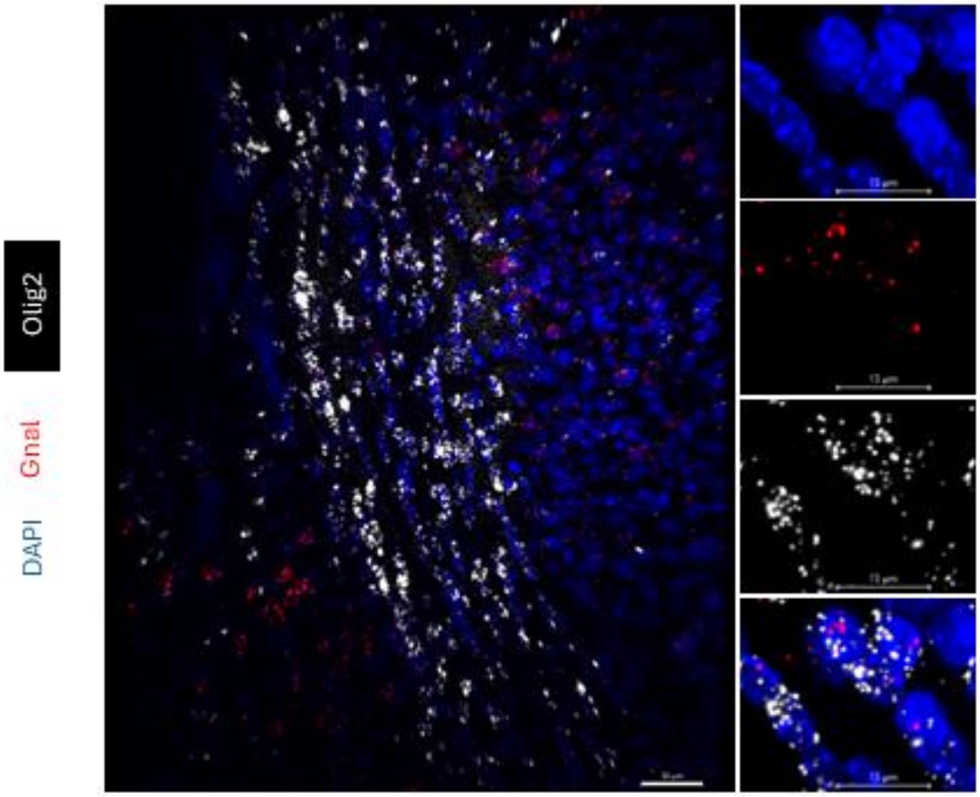

**Supplemental Figure 4: GABAergic Cortical Cells Variably Express *Gnal*. A)** Micrographs of *Gnal* and *Gad1* expression patterns at 10x (top row) and a representative cell at 60x magnification (bottom row). **B)** Quantification of single cell *Gnal* transcript expression average between GABAergic neurons of the cortex. Layer 2/3 displayed a mean of 55.08 transcripts (95% CI 48.09-62.07), while deep cortical layers were found to average 61.18 transcripts (95% CI 54.82-67.54) Expression differences were not statistically significant; $p>0.05$, Unpaired-T test, $t=1.297$. N= 3 mice, n= 50 cells of each classification.

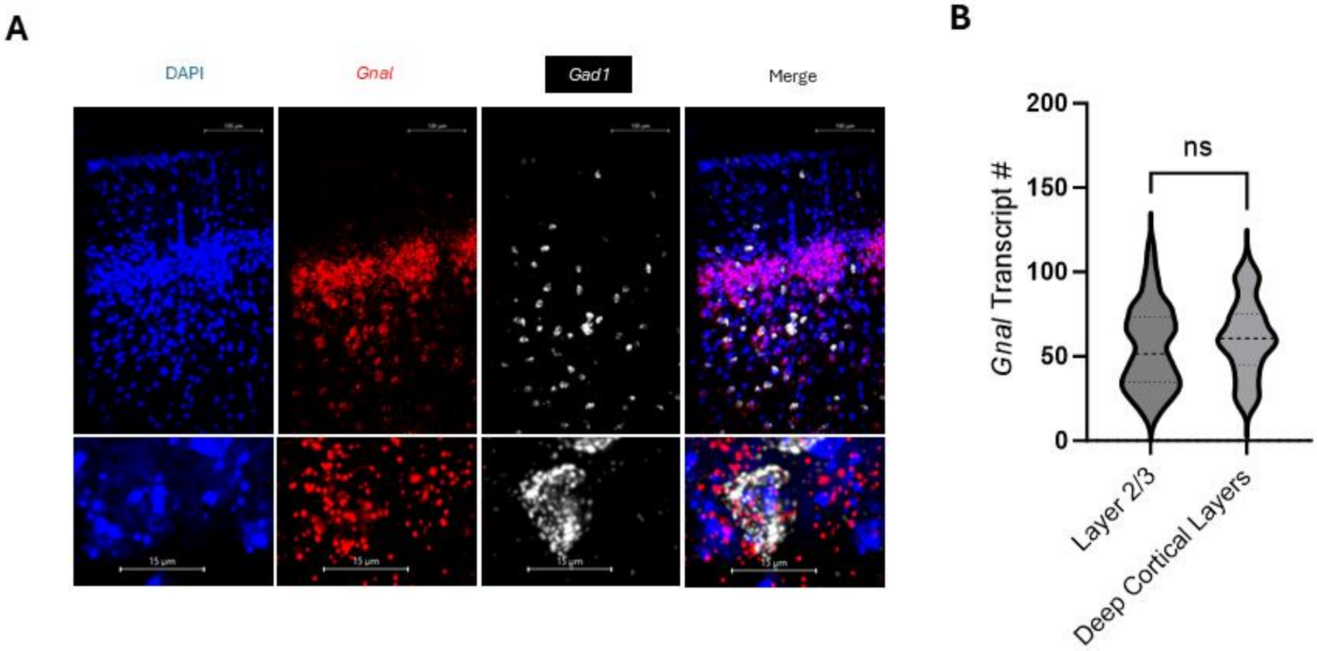


**Supplemental Figure 5: Sparse Cells in the Subiculum Express Substantial *Gnal*. A)** Micrographs of *Gnal* (red) expression in the hippocampus at 10x magnification (left). Micrographs of the subiculum at 60x magnification reveal *Sst* (white) positive cells exhibit low to no *Gnal* expression (right).

**A**

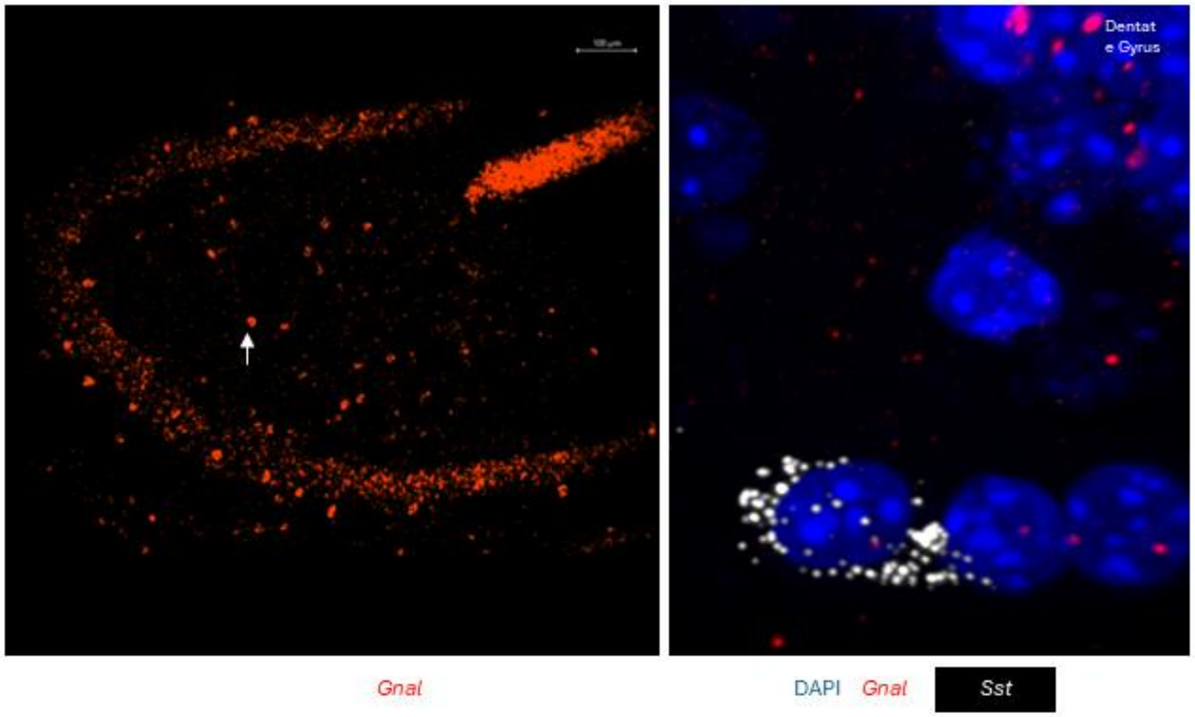


**Supplemental Figure 6: *Gnal* Expression Varies Between Cell Type. A)** Quantification of average *Gnal* transcript expression on a single cell basis. Purkinje cells express the highest average of *Gnal* transcripts on a single cell basis and *Sst* positive striatal interneurons the lowest.

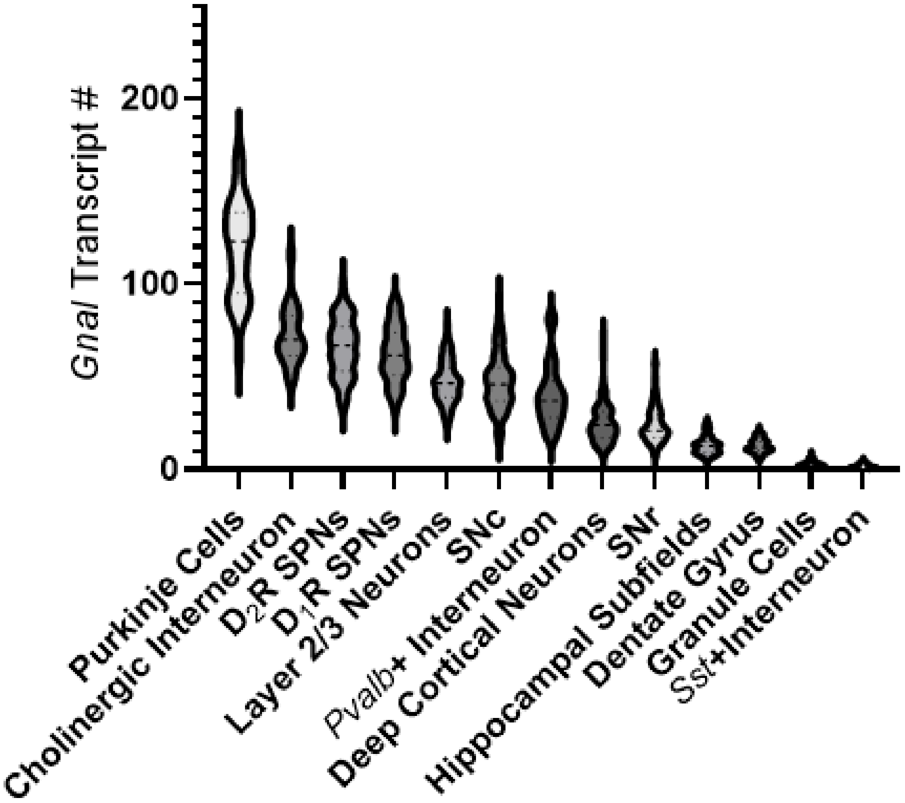

Gnal Transcript #
200
100
0
Purkinje Cells
Cholinergic Interneuron
D2R SPNs
D1R SPNs
Layer 2/3 Neurons
SNc
Pvalb+ Interneuron
Deep Cortical Neurons
SNr
Hippocampal Subfields
Dentate Gyrus
Granule Cells
Sst+Interneuron
